# The effect of varied exercise intensity on antioxidant function, and aortic endothelial cell function and serum lipids in a non-alcoholic fatty liver disease rats

**DOI:** 10.1101/407395

**Authors:** Ling Ruan, Smart Neil.A., Fanghui Li

## Abstract

Exercise and diet may improve cardio-metabolic health in non-alcoholic fatty liver disease, but the optimal exercise prescription remains unclear. We aimed to compare the effects of diet and exercise at different intensities on antioxidant function, and aortic endothelial cell function and serum lipids in a non-alcoholic fatty liver disease rats. Fifty Sprague Dawley rats (180-220g) were randomly divided into two experimental groups and fed either standard rodent chow diet or a high-fat diet. After16 weeks, these animals that received the HFD were randomly separated into a high fat control group or three exercise training groups: HF and low intensity exercise, HF and moderate intensity exercise, HF and incremental intensity exercise, these experimental rats keep sedentary or training for the next 6 weeks. Markers of Aortic Oxidative stress were detected using assay kit. Immunohistochemical analysis was performed to determine the expression level of eNOS and ET-1. Lipid metabolism parameters were detected with an automatic analyzer. Exercise at different intensities improved lipid metabolism, enhanced anti-oxidation function, reduced MDA, increased NO, and improved the expression of eNOS and ET-1 protein levels. Decreased blood lipids were exhibited in all exercise groups. Notably, moderate intensity exercise demonstrated more effect on increasing GSH contents, and decreased the expression of ET-1 protein levels.

## Introduction

Nonalcoholic fatty liver disease (NAFLD) is the hepatic manifestation of metabolic syndrome, and it is the most prevalent liver disease worldwide(1). The prevalence of NAFLD is approximately 30% in the United States and Europe, with a similar prevalence documented in Asian countries (2). It encompasses a spectrum ranging from simple steatosis to fatty liver with hepatocellular injury, termed nonalcoholic steatohepatitis (NASH), fibrosis and cirrhosis(3).

Moreover, the majority of deaths among NAFLD patients are not only associated with liver-related morbidity and mortality but also related to cardiovascular and other complications. A large number of studies have shown that high-fat diets can cause lipid metabolism disturbances, abnormal lipid accumulation, obesity, and NAFLD (4-6). Free fatty acids (FFA) can cause oxidative stress which is a primary cause of intravascular dysfunction, and therefore long-term high-fat diets can inhibit nitric oxide synthase expression in vascular endothelial cells, reduce Nitric oxide (NO) production, resulting in abnormal blood vessels endothelial cell function and vascular endothelial dysfunction. NO is produced via NO synthases, which are a family of enzymes catalyzing the production of NO from L-Arginine. For the purposes of this work we will consider total NO synthase (T-NOS), endothelial NO (e-NOS) and inducible NO (iNOS). T-NOS as the name suggests is the aggregate NO synthase circulating at any particular time, while e-NOS is the endothelial NOS generated in blood vessels and is involved with regulating vascular function. i-NOS is inducible NOS which is usually raised in an oxidative environment. As NO expression is altered with endothelial dysfunction, which in turn is associated with NAFLD, finding an effective management solution is therefore a current research priority.

Previous experiments have shown aerobic exercise can improve lipid metabolism, oxidative stress(7, 8) and vascular endothelial function(9). Several pharmacological and non-pharmacological strategies have been proposed to relieve NAFLD-associated deleterious alterations (10). Among non-pharmacological approaches, physical exercise-mediated multi-systemic adaptations can promote crosstalk between organs and orchestrate pro-metabolic effects known to mitigate metabolism-related disorders such as NAFLD(11). Recently, Keating SE et al. examined the efficacy of commonly prescribed exercise dose and intensity for reducing liver fat and visceral adipose tissue in an animal experimental model of NAFLD, but no significant differences were found between the dose or intensity of the exercise regimen and reductions in liver fat or visceral adipose tissue (12). Paradoxically, it has been shown that vigorous and moderate exercise were equally effective in reducing intra-hepatic triglyceride content, but body weight, body fat, waist circumference, and blood pressure with vigorous-moderate intensity exercise was lower than the moderate intensity group(13, 14). Similarly, Tsunoda K *et al*. showed vigorous-intensity was more effective than moderate-low intensity exercise and moderate-high intensity protocols in preventing nonalcoholic fatty liver from progressing to NASH (14). Two systematic reviews of published studies of NAFLD patients participating in aerobic exercise programs showed that liver fat was significantly reduced, but the optimal exercise intensity is undetermined(15, 16), although a growing number of prospective data shows the effects on different types of exercise on NAFLD(17). Collectively, these previous findings suggest that the intensity of exercise, rather than the volume or duration, may play a critical role in magnifying the protective effects against NAFLD (18).

The relationship between non-alcoholic fatty liver disease and aortic endothelial function is poorly understood. It is unclear whether exercise could affect aortic endothelial function in a dietary-induced rat model of NAFLD. Moreover, different exercise intensities may produce varying effects on endothelial function in rat NAFLD model. Therefore, this study compared the effect of different exercise intensities, on markers of aortic endothelial function in a high fat diet-induced NAFLD rat model.

## Materials and Methods

### Animals

Male Sprague–Dawley (SD) rats (180-220g) were purchased from the Guangdong Medical Laboratory Animal Center (GDMLAC) (Guangzhou, China). Rats were raised in a SPF environment (23 ± 1°C,humidity 60-70%, 12h light/dark cycle), in the Laboratory Animal Center. SD rats (n=50) were randomly divided into two experimental groups and fed either standard rodent chow diet control group (CON; n=10), or a high-fat diet (HFD; n=40), after16 weeks, animals that received the HFD were randomly separated into a sedentary control high fat group (HFC; n=10) or three exercise training groups: HF and low intensity exercise (LE; n = 10), HF and moderate intensity exercise (ME; n = 10), HF and incremental intensity exercise (IE; n =10). For the next 6 weeks, CON group libitum feeding with standard rodent chow diet and remained sedentary, HF group libitum feeding with high fat diet and remained sedentary. LE, ME and IE groups were fed with high fat diet and trained with different exercise training intensities. At the end of 6 weeks treatment, rats were sacrificed after fasting overnight. Blood sample, aorta samples, and liver samples were harvested for analysis. All the procedures were performed in compliance with the Institute’s guidelines and with the Guide for the Care and Use of Laboratory Animals published by the US National Institutes of Health (NIH Publication No. 85-23, revised 1996). The study was approved by the institutional animal care committee of GDMLAC.

### Composition of the high fat diet

The high–fat diet (HF, GDMLAC) contained several compounds that provide energy (5% sucrose, 18% lard, 15% egg yolk powder, 0.5% sodium cholate and 1% cholesterol add to the 60.5% basic standard rodent chow diet; GDMLAC).

### Exercise experimental protocol

All animals were familiarized with treadmill running (DSPT202, Qianjiang Technology Company, Hangzhou, China) at 0–15 m/min, 10–20 min per day for six consecutive days. An electrified grid (0.6–mA intensity) was placed behind the belt of the treadmill to induce running. The rats that failed to run regularly were excluded from the training protocol. The exercise program involved 60 min/day, 5 days per week, totally 6 weeks. The daily training intensity program for each group respectively: LE group: 15 m/min, ME group: 20 m/min, and IE group consisted of running 10 minutes at 15 m/min, followed by a gradual increase in intensity at 20 m/min for 30 min, and increase in intensity to 27 m/min for 20 min on a motor–driven treadmill.

### Outcome Measures

The primary outcome measures were the markers of oxidative stress in the aorta; T-NOS and i-NOS. SOD, MDA, CAT, GST and T–AOC.

The secondary outcome measures were; presence of aortic enothelin-1 and eNOS, body mass and liver mass and lipids.

We also confirmed the existence of NAFLD (liver histology) by hematoxylin and eosin (HE) staining of embedded liver tissue samples.

### Markers of Aortic Oxidative stress

The aorta was separated from the ice plate and then put it in liquid nitrogen preserve the sample for testing. Prepared fresh aorta samples were ground in saline solution to make 10% aorta homogenates, followed by centrifugation for 20 min at 4°C. The resulting supernatant was collected using specific kits according to the manufacturer’s instructions. NOS (T-NOS and iNOS) were detected using an assay kit (Colorimetric method), (Nanjing Jiancheng Corp., Nanjing, China), Nitric Oxide (NO) assay kit (Nitrate reductase method).

MDA contents in the aorta were quantified using a lipid peroxidation MDA assay kit (TBA method) (Beyotime Institute of Biotechnology, Jiangsu, China) according to the manufacturer’s protocol. CAT activity assay kit (Visible light method), SOD, T–AOC activity and GST contents were determined using a reagent kit (Colorimetric method), (Nanjing Jiancheng Corp., Nanjing, China).

### Endothelin-1 and Nitric oxide synthase

Endothelin-1 (ET-1) and endothelial nitric oxide synthase (eNOS) were measured by immune-histochemical analysis. After the abdominal cavity was opened, the aorta was quickly and completely separated, the fixed aorta was embedded in paraffin, sliced into 5-μm-thick sections, and mounted on glass slides. The immunohistochemistry was performed with a PowerVision two-step immunohistochemistry detection kit. ET-1, and eNOS antibodies were obtained from (Bioss Biotechnology Co., Ltd. Beijing, China). Samples were observed through JVC3-CCD camera (Nikon Corp., Tokyo, Japan), and Image-Pro Plus image (Media Cybernetics Corp., USA) processing software system was used for image acquisition and analysis. The brown granules visible in the cytoplasm or nucleus were considered positive expression of aortic endothelial cells. The number of positive cells per section was counted in 10 random fields (400x magnification), and the percentage of positive cells (positive cells/total cells × 100%) was calculated. Three non-consecutive sections were selected from each specimen and those indices were averaged.

### Lipids

Blood samples were collected from the abdominal aorta and centrifuged at 3000 rpm for 15 min, and then serum was collected. The serum triglyceride (TG) levels (mmol/l), total cholesterol (TC) levels (mmol/l), low–density lipoprotein cholesterol (LDL–c) levels (mmol/l), high-density lipoprotein cholesterol (HDL–c) levels (mmol/l), were detected with an automatic analyzer (Toshiba AccuteTBA-40FR, Toshiba Corporation, Tokyo, Japan).

### Characterization of non-alcoholic fatty liver disease

To characterize NAFLD paraffin–embedded liver tissue samples were cut into 5-μm thick sections for hematoxylin and eosin (HE) staining and the sections were then examined by light microscopy. Rat livers were fixed with 10% formalin, cut into 10–μm sections with hematoxylin and eosin and then at least three randomly selected liver sections images were digitally captured (400x magnification), from each group were examined and photographed with a Nikon Eclipse Ci light microscope (Nikon Corp., Tokyo, Japan).

### Statistical analysis

Statistical analysis was carried out using the Statistical Package for the Social Science (SPSS version 20.0, IBM Corp., USA). Data are presented as mean ± standard deviation. Graph Pad Prism (version 5.0; Graph Pad Prism Software, La Jolla, CA, USA). One–way ANOVA and Tukey’s significant difference post-hoc analysis were conducted. P value of ≤ 0.05 denoted a statistically significant difference.

## Results

### Effect of exercise training intensity on aortic NOS and NO activity

Table 1 shows that aortic T-NOS activity was higher in the CON versus HFC (*P*<0.01) and IE (*P*<0.05) group, however only the low intensity (LE) group showed a significant elevation compared to HFC (*P*<0.05). i-NOS activity was higher in the HFC and all exercise groups versus CON group (*P* <0.01), however only the low LE and ME groups showed a significant reduction compared to HFC (*P*<0.01), meanwhile the LE and ME groups showed a significant reduction compared to the IE group (*P* <0.01).

**Table 1.**
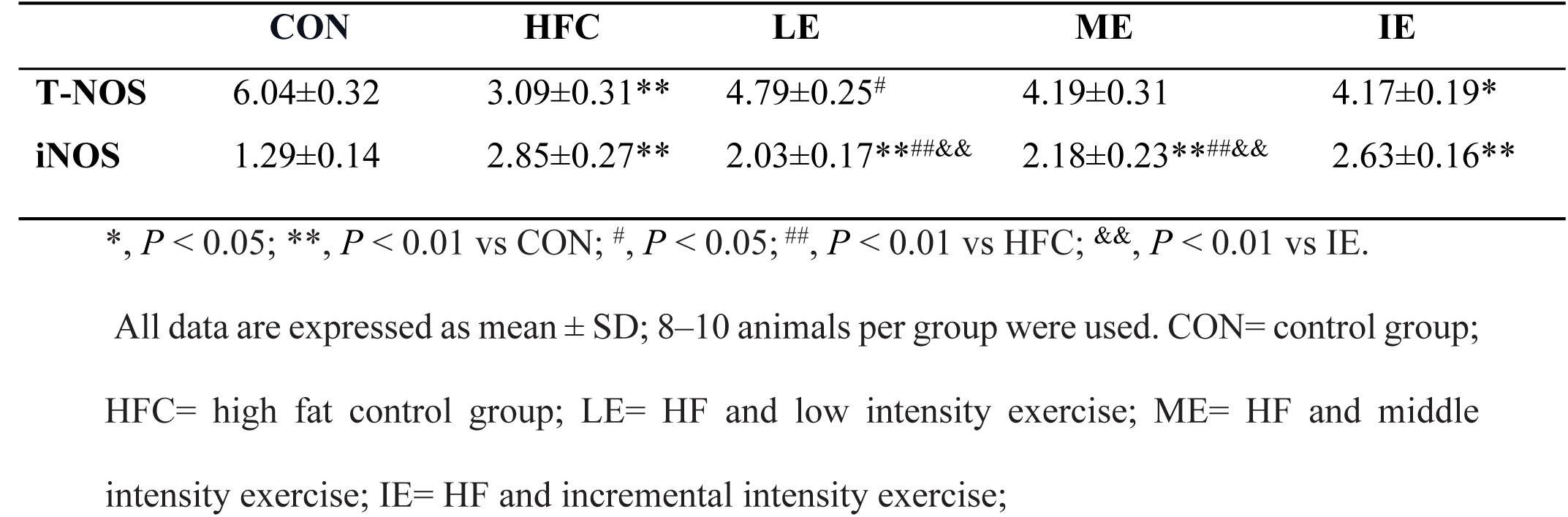
The Change of Aortic Nitric Oxide Synthase (NOS) activity in Each Group(U/mg prot)

Table 2 shows that NO content in the aorta was higher in the CON versus HFC (*P*<0.01) and all exercise (*P*<0.01) groups, all exercise groups showed a significant elevation compared to HFC (*P*<0.01).

**Table 2.**
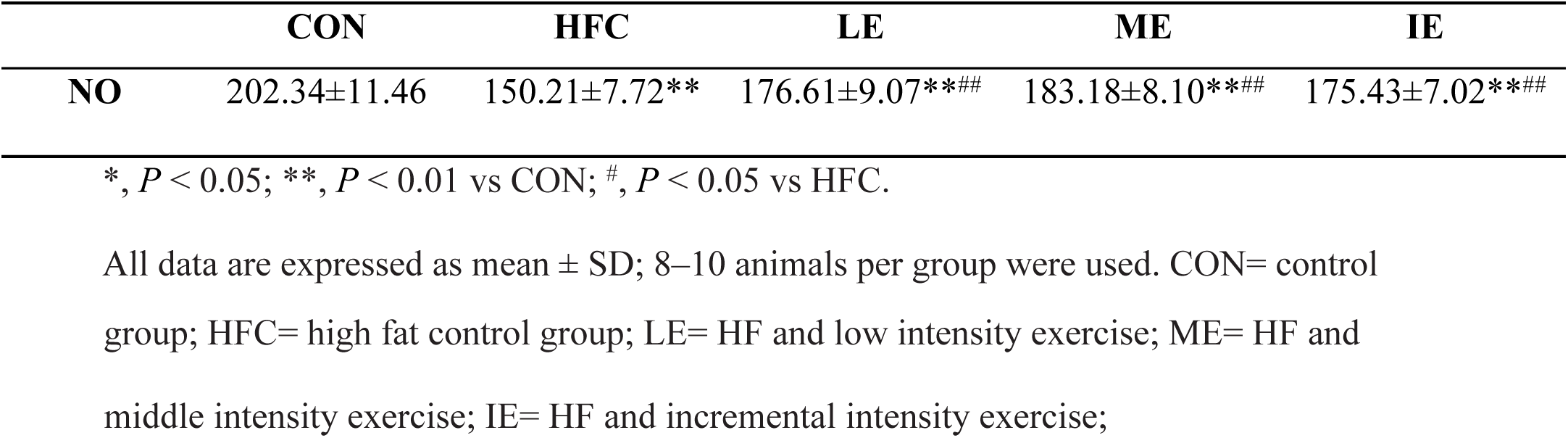
The Change of Aortic NO in Each Group(U/mg prot)

### Effect of different exercise training intensities on aortic endothelial cell oxidative stress

As shown in Fig 1, compared with the CON group, CAT, GSH, and T–AOC indicated a significant reduction in the HFC group (*P*<0.01; Figs 1C, 1D and 1E). After 6 weeks exercise training, SOD, CAT, and T–AOC in LE, ME, and IE group were significantly higher compared to the HFC group (Figs 1A, 1C and 1E), whereas the MDA levels were decreased compared to the HFC group (*P*<0.01; Fig 1B). In addition, LE and ME groups exhibited significantly increased GSH compared to the HFC group (*P*<0.05 and *P*<0.01 respectively; Fig 1D).

**Figure 1.** Effects of exercise training on markers of oxidative status in the aortic endothelial cell. SOD (A), MDA (B), CAT (C), GST (D), and T–AOC (E). **, *P* < 0.01 compared with CON group; ^#^, *P* < 0.05; ^##^, *P* < 0.01 compared with HFC group; All data are expressed as mean ± SD; 8–10 animals per group were used. CON= control group; HFC= high fat control group; LE= HF and low intensity exercise; ME= HF and middle intensity exercise; IE= HF and incremental intensity exercise;

### Effect of different exercise training intensities on eNOS and ET-1 expression in the aorta

Expression of eNOS protein levels in the aorta were significantly lower in HFC group than in CON group (*P*<0.05), whereas there was no difference between the three exercise groups (Table 3 and Fig 2). The expression of ET-1 protein levels were significantly higher in the HFC, LE, and IE groups compared to the CON group. The ET-1 levels were significantly decreased by moderate intensity exercise training (*P*<0.01). (Table 4 and Fig 3).

**Table 3.**
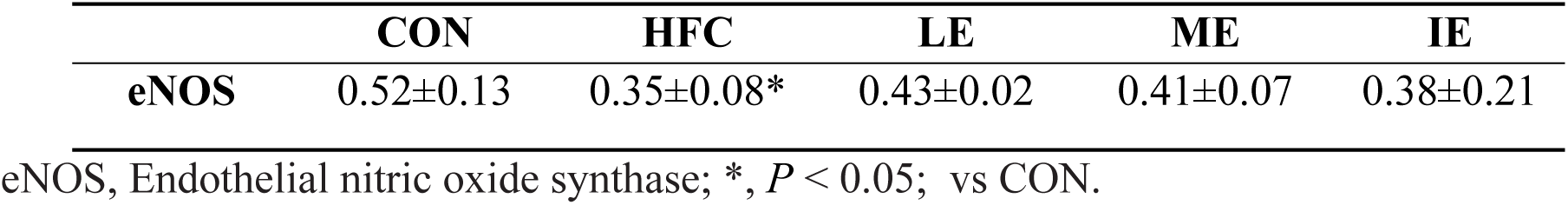
The Change of eNOS expression in Each Group

**Table 4.** The Change of ET-1 expression in Each Group

|  | CON | HFC | LE | ME | IE |
| --- | --- | --- | --- | --- | --- |
| **ET-1** | 0.25±0.03 | 0.39±0.04** | 0.33±0.05* | 0.29±0.02<sup>##</sup> | 0.35±0.09** |

ET-1, Endothelin-1; \*, $P < 0.05$ ; \*\*, $p < 0.01$ vs CON; <sup>##</sup>, $P < 0.01$ vs HFC.

**Figure 2.** Effects of exercise training on aortic eNOS expression.

**Figure 3.** Effects of exercise training on aortic ET-1 expression.

All data are expressed as mean ± SD; 8–10 animals per group were used. CON= control group; HFC= high fat control group; LE= HF and low intensity exercise; ME= HF and middle intensity exercise; IE= HF and incremental intensity exercise;

All data are expressed as mean ± SD; 8–10 animals per group were used. CON= control group; HFC= high fat control group; LE= HF and low intensity exercise; ME= HF and middle intensity exercise; IE= HF and incremental intensity exercise;

All data are expressed as mean ± SD; 8–10 animals per group were used.

CON= control group (A); HFC= high fat control group (B); LE= HF and low intensity exercise (C); ME= HF and middle intensity exercise (D); IE= HF and incremental intensity exercise (E).

All data are expressed as mean ± SD; 8–10 animals per group were used. CON= control group (A); HFC= high fat control group (B); LE= HF and low intensity exercise (C); ME= HF and middle intensity exercise (D); IE= HF and incremental intensity exercise (E).

### Effect of different exercise training intensities on body mass and liver mass

As shown in Fig 4, after 6 weeks treatment, body mass of the rats in each group were not significantly different (Fig 4A). Liver mass were lower in the ME group than in the HFC group (*P*<0.05), otherwise there was no significant difference among the exercise groups (Fig 4B).

**Figure 4.** Effects of exercise training on body mass, liver mass. Body mass (A), liver mass (B) of each group. *, *P*< 0.05 compared with CON group; ^#^, *P* < 0.05 compared with HFC group; all data are expressed as mean ± SD; 8-10 animals per group were used. CON= control group; HFC= high fat control group; LE= HF and low intensity exercise; ME= HF and middle intensity exercise; IE= HF and incremental intensity exercise;

### Effect of different exercise training intensities on lipid metabolism disorders and liver histology

As shown in Fig 5, liver histology was evaluated by H&E staining. CON group rats tissue exhibited well-arranged hepatic cords, cells with round and central nuclei, a lobular structure and an array of wheel–shaped cells along the centrilobular vein. However, in the HFC group lipid droplets were observed in the liver sections (Fig 5B). Lipid droplet volumes and quantities were reduced with different exercise intensities (Figs 5C, 5D, and 5E).

**Figure 5.** The Optical Microscope Image of H&E staining in Rat Liver Tissue(400×)

CON= control group (A); HFC= high fat control group (B); LE= HF and low intensity exercise (C); ME= HF and middle intensity exercise (D); IE= HF and incremental intensity exercise (E);

As shown in Table 5, the serum TC, TG, LDL-c and FFA were lowest in the CON group, but TC, TG and LDL-c were significantly decreased in the LE, ME and IE groups compared with the HFC group. No difference in serum HDL-c was observed between groups. Notably, TG, TC, and LDL were not significantly different between the three exercise groups.

**Table 5.**
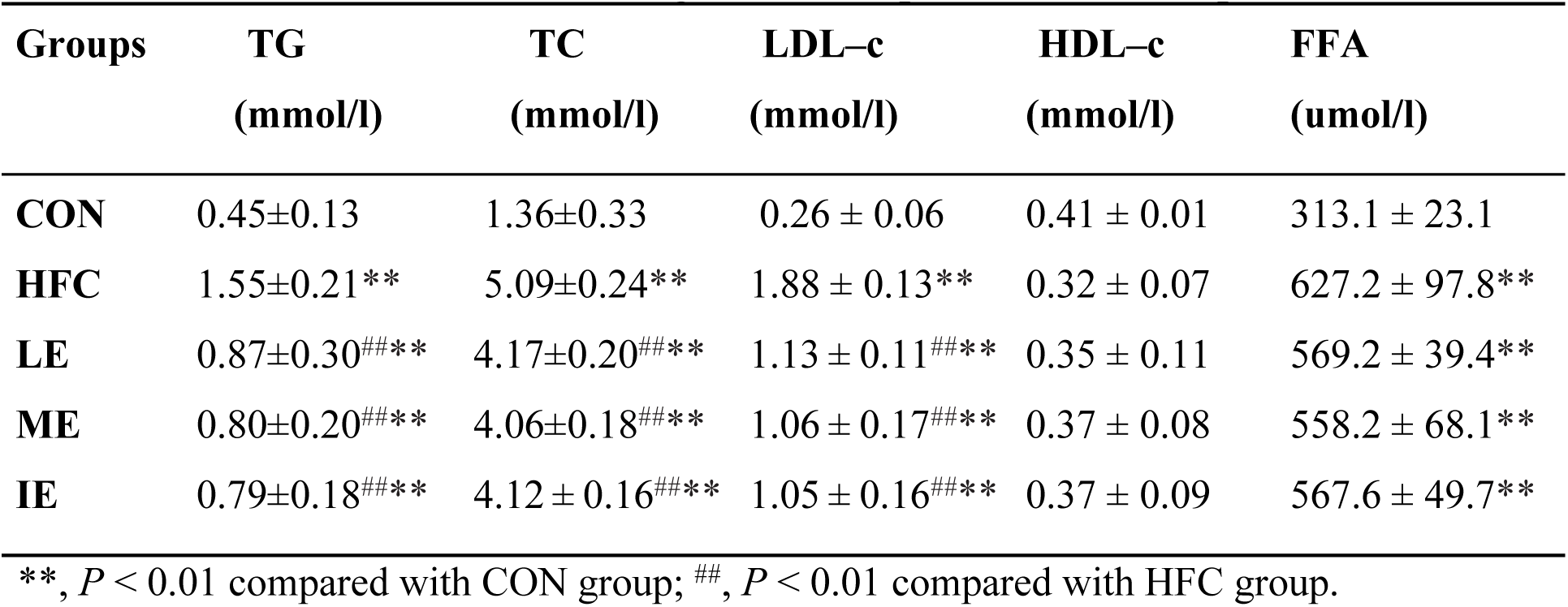
The Change of Blood lipids in Each Group

All data are expressed as mean ± SD; 8–10 animals per group were used. CON= control group; HFC= high fat control group; LE= HF and low intensity exercise; ME= HF and middle intensity exercise; IE= HF and incremental intensity exercise;

## Discussion

The aim of our study was to investigate the effects of varying exercise intensities on aortic endothelial function in high fat diet-induced NAFLD rats. We can confirm that high fat diet induced vascular endothelial dysfunction in NAFLD rats. We found that exercise enhanced anti-oxidation function and improved some markers of aortic endothelial cell function. T-NOS activity appeared to respond best to low intensity exercise, i-NOS activity was higher in only the low intensity and moderate intensity groups. Moderate intensity exercise demonstrated the greatest effect on decreasing the expression of the potent vasoconstrictor ET-1 levels, whereas GSH was raised in this group. Decreased blood lipids were exhibited in all exercise groups.

### The impact of different exercise training intensities on antioxidant function of aortic endothelial cells

Our study showed that CAT, GSH, and T–AOC was significantly reduced in the HFC group, compared with the CON group, confirming the desired effect of the HF diet inducement of a NAFLD rat model. After 6 weeks exercise training SOD, CAT, and T–AOC were significantly increased, conversly, MDA was significantly reduced in all exercise groups. Furthermore, low intensity and moderate intensity exercise increased GSH.

Previous work has shown that high fat diets increase lipid peroxidation and destroy the balance of the oxidative and antioxidative systems (19). Moreover, oxidative stress and increased ROS production are the primary cause of dysfunction in aortic endothelial cells (20, 21). The relationship between exercise and oxidative stress is extremely complex, depending on the mode, intensity, and duration of exercise. Pingitore A *et al*. noted that regular moderate training in humans appears beneficial for oxidative stress and health, conversely, acute exercise leads to increased oxidative stress (22), presumably as their a period of adaptation that is missing from acute exercise training. Pereira *et al.* also showed high-intensity exercise may induce oxidative stress (23). Li *et al*. reported that SOD activity and GSH were significantly raised after rats were exercised at medium intensity (24). Radak *et al*. also indicated that moderate exercise significantly increased the activity of antioxidant enzymes(25). However Lu *et al*. reported that high intensity exercise was superior to the moderate intensity in attenuating oxidative stress and improving glucolipid metabolism in post-MI rat myocardium (26). Jamurtas *et al*. also found high intensity to be superior to moderate intensity for reducing oxidative stress in healthy male humans(27). It therefore remains unclear which exercise intensity is optimal for improving anti-oxidant function.

In our study, we elucidated that all exercise training intensities, especially for low intensity and moderate intensity, enhanced antioxidant enzyme activity and suppressed NAFLD rats’ oxidative stress. We speculated, and our data supported, the notion that that incremental exercise may increase reactive oxygen species(ROS) production during incremental exercise leading to the oxidation of protein, lipids or nucleic acid (28). The production of ROS during exercise is also accompanied by a reduction of antioxidant capacity (29). However, our data lack measures of ROS production and other related oxidative stress markers. So further studies of the molecular mechanisms involved in anti-oxidation may be indicated.

### The impact of different exercise training intensities on endocrine function of aortic endothelial cells

Our results show that the expression of eNOS and ET-1 levels were significantly reduced in the moderate intensity group. The activity of i-NOS was significantly increased in the low and moderate intensity groups, compared to the incremental (high) intensity group, this may be due to greater oxidative stress in the latter group, leading to iNOS consumption. ET-1 was reduced in the moderate intensity group. Together these results suggest, exercise a low or moderate intensity elicited greatest improvements in ET-1 and Nitric Oxide profile, implying that low to moderate intensity exercise is optimal for protection of endothelial cell function.

Other work has shown exercise training, at symptom-limited intensity, improves arterial endothelial cell function in people with heart disease (30). Several previous studies have shown low and moderate intensity exercise to have a positive effect on aortic endothelial cell function of rats (31, 32). Shaodong *et al*. indicated that aerobic exercise, of unknown intensity, decreases the production of lipid oxidation products, and thus prevents damage to endothelial cells in rats with dyslipidemia(33). Furthermore, a previous experiment demonstrated that moderate intensity exercise could reduce expression of ET-1 levels, induced by aortic injury in mice (34). Wang *et al*. and Archana *et al*. found that moderate intensity exercise is optimal for raising serum NO (7, 35). However, Morishima T *et al*. indicated endothelial function was was maintained by conducting high-intensity resistance exercise (36). A recent meta-analysis showed that high intensity training seems to have a superior effect on the improvement of endothelial function compared with moderate exercise in cardiac patients (37). All these results demonstrate exercise improves the function of aortic endothelial cells, however, the optimal exercise intensity remains unclear.

We speculate that moderate exercise demonstrated improved aortic endothelial cell function, is underlined by the following: (i) Moderate exercise improves lipid metabolism, promoting fat mobilization and lipid energy catabolism [17](38). (ii) Moderate intensity exercise has a beneficial anti-oxidative effect [38]. (iii) The reduced ET-1 and increased NO expression were significant in the moderate intensity exercise group, ultimately improving aortic endothelial cell function (39, 40). We do however concede that results of other work are conflicting.

### The impact of different exercise training intensities on lipid metabolism disorders

We demonstrated that three different exercise training intensities were equally effective in alleviating dyslipidemia as well as hepatic damage in a diet induced rat NAFLD model. These results suggested that the therapeutic effect of exercise training in dyslipidemia and hepatic damage is unrelated to exercise intensity. This notion is supported by meta-analytic work (41). Moreover, our study did not find any improvement in body mass or HDL-c in any group. However, it should be noted that liver mass was significantly decreased in the moderate intensity group.

Exercise plays an important role in improving lipid metabolism disorders, and is increasingly seen as an adjunctive therapy for the prevention and treatment of NAFLD (16, 17, 42). Machado MV *et al.* showed that exercise intensity would be more effective in improving metabolic parameters than frequency or duration (43). A retrospective study indicated that moderate and vigorous intensity physical activity yielded similar health benefits to low, in terms of the measured body adiposity and serum TG (38). Two studies showed that vigorous-intensity interval training and continuous moderate-intensity exercise have the same effect on lowering the serum FFAs, TG of NAFLD in animals (17, 44). Suk M *et al*. also indicated that high intensity exercise improved lipid metabolism in the liver of rats(45). Fisher *et al.* also did not find any improvements in body weights and HDL-c between groups of differing exercise intensity (46). However, Khammassi M *et al*.showed that high-intensity interval training may be particularly useful in overweight/obese youth to improve body composition and lipid profile (47).

## Conclusions

The role and underlying mechanism of exercise training in NAFLD related aortic endothelial cell function remain poorly understood. Exercise at different intensities, enhanced anti-oxidation function, increased NO contents, decreased MDA. Moreover, reduced the expression of aortic eNOS and ET-1 protein levels, improved lipid metabolism. Notably, moderate intensity exercise demonstrated more effect on decreasing the expression of ET-1 protein levels, and GSH.

## Supporting information

**S1 Fig 1. Effects of exercise training on markers of oxidative status in the aortic endothelial cell.**

**S2 Fig 2. Effects of exercise training on aortic eNOS expression.**

**S3 Fig 3. Effects of exercise training on aortic ET-1 expression.**

**S4 Figure 4. Effects of exercise training on body mass, liver mass**

**S5 Fig 5. The Optical Microscope Image of H&E staining in Rat Liver Tissue(400×)**

## Abbreviations

NAFLD: non-alcoholic fatty liver disease
NASH: nonalcoholic steatohepatitis
HFD: high fat diet
HFC: high fat sedentary control
LE: low intensity exersise
ME: moderate intensity exercise
IE: incremental internsity exercise
TG: Triglyceride
TC: total cholesterol
HDL-c: HDL-cholesterol
LDL-c: LDL-cholesterol
FFA: free fatty acid
SOD: superoxide dismutase;
MDA: malondialdehyde;
CAT: catalase;
GST: glutathione;
T–AOC: total antioxidant capacity
ET-1: endothelin1
T-NOS: Total Nitric Oxide Synthase
eNOS: endothelial nitric oxide synthase
iNOS: inducible nitric oxide synthase
NO: Nitric Oxide

